# Data Descriptor: Proteomic profiling across breast cancer cell lines and models

**DOI:** 10.1101/2020.12.15.422823

**Authors:** Marian Kalocsay, Matthew J. Berberich, Robert A. Everley, Maulik K. Nariya, Mirra Chung, Ben Gaudio, Chiara Victor, Gary A. Bradshaw, Marc Hafner, Peter K. Sorger, Caitlin E. Mills, Kartik Subramanian

## Abstract

We performed quantitative proteomics on 61 human-derived breast cancer cell lines to a depth of ~13,000 proteins. The resulting high-throughput datasets were assessed for quality and reproducibility. We used the datasets to identify and characterize the subtypes of breast cancer and showed that they conform to known transcriptional subtypes, revealing that molecular subtypes are preserved even in under-sampled protein feature sets. All datasets are freely available as public resources on the LINCS portal. We anticipate that these datasets, either in isolation or in combination with complimentary measurements such as genomics, transcriptomics and phosphoproteomics, can be mined for the purpose of predicting drug response, informing cell line specific context in models of signalling pathways, and identifying markers of sensitivity or resistance to therapeutics.

## Background & Summary

Targeted therapy relies on the identification of actionable changes in signal transduction, proliferation or cell death pathways that are drivers of transformed states. In some cases these changes are associated with a recurrent mutation or overexpression of an oncogene. In other cases, the causes of differences in drug sensitivity are less well understood. Some breast cancer subtypes are particularly responsive to targeted therapy owing to high expression of one or more of the estrogen (ER), progesterone (PR), or HER2 receptors. Moreover, the presence or absence of these receptors, which is most commonly measured by immunohistochemistry (IHC), defines clinical breast cancer subtype and mode of first line therapy (expression of ER and/or PR defines the hormone receptor (HR) positive subtype and over-expression of HER2/ERBB2 defines the HER2 positive subtype). The third breast cancer subtype, triple negative breast cancer (TNBC), lacks high levels of ER, PR and HER2 expression, is genetically heterogeneous and is the least effectively treated^1^. It is therefore of considerable interest to identify recurrent changes in TNBCs that might be targeted to treat disease. Breast cancers can also be classified into molecular subtypes based on gene expression signatures. These include the luminal A/B and basal designations that generally encompass HR positive and TNBC disease respectively, with HER2 enriched cancers comprising a separate molecular subtype^2–4^.

Multiple studies have been performed in which panels of TNBC cell lines are subjected to transcript profiling to identify differences among them^5,6^. However, transcript levels do not necessarily correlate with the abundance of proteins, which are the ultimate targets of small molecule and antibody therapies^7^. Moreover, in some tumor types, the effects of copy number alterations extend to mRNA abundance without necessarily propagating to changes in protein abundance^8^. It is therefore valuable to measure the levels of proteins across panels of cell lines as a means to identify changes in cell state. Of particular interest are changes that might individually or in combination determine sensitivity to new or existing drugs for the treatment of breast cancer. It has been shown that computational models of drug sensitivity that are trained using protein expression data can complement or even outperform models trained on transcript or genomic data alone^9–11^. Thus, a standardized dataset on protein expression levels in breast cancer cell lines of all three clinical subtypes is expected to constitute a valuable resource for drug discovery and development of predictive biomarkers.

In this data descriptor we describe systematic profiling of 60 widely used breast cancer cell lines using Tandem Mass Tag (TMT) liquid chromatography mass spectrometry (LC/MS). TMT LC/MS is a method for labelling multiple samples (up to 11 in the current work) with mass tags and then analysing them in a single run of the mass spectrometer, thereby enabling direct comparison of protein levels. In this data descriptor we provide a technical summary of the TMT based mass spectrometry approach and the resulting data. Quality metrics used to asses both technical and biological validity are explained and we discuss how the resulting data can be leveraged to characterize preclinical (cell line) models of breast cancer, generate testable hypotheses of resistance to therapy and discover novel biological insight.

## Methods

### Culture Conditions

All cell lines used in this study were of human female origin and derived from breast cancers except the 184A1, 184B5, MCF 10A, MCF12A and HME1 cell lines which were derived from non-malignant human breast epithelia. Cell lines were maintained, free of mycoplasma, in their recommended growth conditions (as listed below), and were identity-validated by STR profiling^12^.

### Mass Spectrometry

A schematic description of our mass spectrometry workflow is shown in **Figure 1**. Data were collected in 8 separate mass spectrometry runs. Because data collection spanned many months and instrumentation and protocols improved over this period, methods differed slightly between batches (runs 1-4 and runs 5-8) as described below.

**Figure 1.**
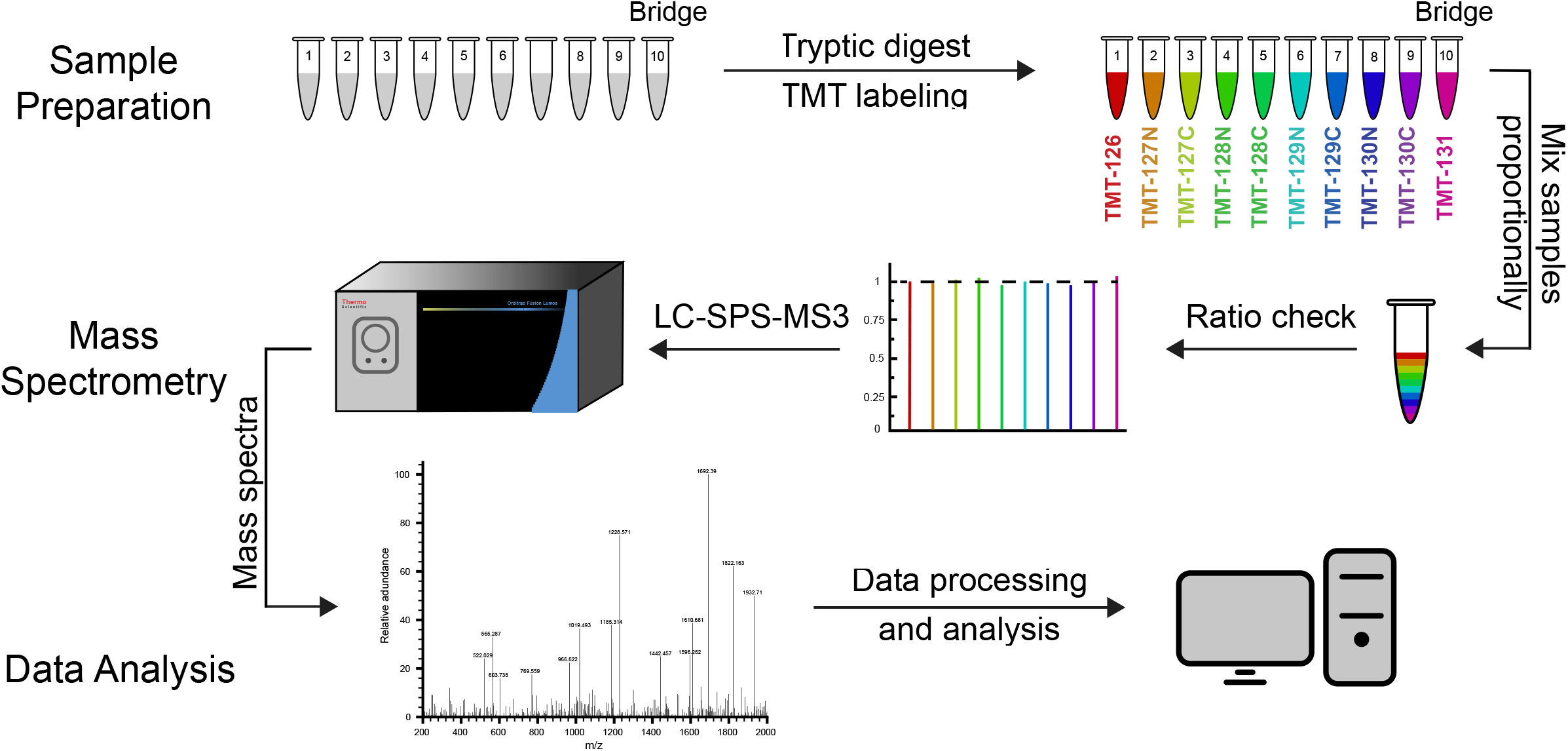
Mass spectrometry workflow. Cell pellets were lysed and either 65 or 150 ug of protein of each sample was labelled using a TMT Mass Tag Labelling Kit (see methods for details of differences between runs). TMT labelled samples were pooled into a single multiplexed sample and a ratio check was performed to ensure that an equal amount of each TMT label was included. The samples were injected into an Orbitrap Fusion Lumos Tribrid mass spectrometer, and TMT quantification was performed in the Orbitrap using SPS-MS3. Assignment of MS/MS spectra was performed using Sequest.

#### Sample Collection

Cells grown in their recommended growth medium to ~60% confluence were rinsed twice with phosphate-buffered saline (PBS) and then gently scraped from 15 cm dishes in PBS supplemented with protease and phosphatase inhibitors (Halt™ Protease and Phosphatase Inhibitor Single-Use Cocktail, EDTA Free, ThermoFisher, Catalog Number 78441) followed by centrifugation at 300 g for 5 minutes at 4°C. The supernatant was discarded and pellets were snap frozen in liquid nitrogen and stored at −80° C.

#### Protein Solubilisation and Digestion

Cell pellets were solubilized in lysis buffer (2% SDS, 150 mM NaCl, 50 mM Tris pH 7.4) supplemented with Halt™ Protease and Phosphatase Inhibitor Single-Use Cocktail, EDTA Free (ThermoFisher, Catalog Number 78443) with a hand-held tissue homogenizer. Disulfide reduction was performed by adding dithiothreitol (DTT) to a final concentration of 5 mM and heating to 37°C for 1 hour, followed by alkylation of cysteine residues with iodoacetamide at a final concentration of 15 mM and incubation at room temperature in the dark for 30 minutes. Protein concentration was determined using a Micro BCA™ Protein Assay Kit (ThermoFisher, Catalog Number 23235) following the manufacturer’s protocol. Detergent was removed by methanol/chloroform protein precipitation as follows: ice cold methanol (3 parts lysis buffer volume), chloroform (2 parts lysis buffer volume) and water (2.5 parts lysis buffer volume) were added sequentially with vortexing after each addition followed by centrifugation at 4000 x g for 10 min. The top layer was aspirated while taking care not to disrupt the interface. Ice cold methanol (3 parts lysis buffer) was added, the samples were vortexed, centrifuged (4000 x g, 10 min.), and the supernatant aspirated leaving behind the protein pellet; this methanol wash procedure was repeated a total of three times^13^.

Precipitates were solubilized in freshly prepared 8 M urea in 200 mM EPPS, pH 8.5. 60 μg of solubilized total protein from each sample was then used for TMT labelling. Following a 10 min incubation at 37°C, the urea concentration was reduced by dilution with 200 mM EPPS, pH 8.5 to 4 M (runs 1-4) or 2 M (runs 5-8) final urea concentration and digestion was then performed by overnight incubation at room temperature in the presence of Lys-C protease (Wako, Catalog Number 129-02541) at an enzyme-to-substrate ratio of 1:75. Following further dilution of the sample with 200 mM EPPS to a final urea concentration of 1.6 M (runs 1-4) or 0.5 M (runs 5-8), digestion was continued by incubation of the sample at 37°C for 6 hours with trypsin (Promega, Catalog Number V5113) at an enzyme to substrate ratio of 1:75. Aliquots corresponding to 65 μg per sample (runs 1-4) or 150 μg per sample (runs 5-8) were withdrawn for TMT labelling.

#### Digest Check

Aliquots equivalent to 5-10 μg from two samples were pooled, desalted and peptides purified by reverse phase chromatography on stage tips^14^ (described below). The peptides were then dried and resuspended in 3% acetonitrile, 5% formic acid (FA) to a final concentration of ~2 μg/μl. The missed cleavage rate was measured by LC-MS/MS to evaluate the quality of the digest; a result under 15% of potential cleavage sites remaining uncleaved was deemed sufficient to proceed with labelling.

#### TMT Labelling, Ratio Check and HPLC Fractionation

Equal amounts of protein were removed from each sample and labelled using a TMT10plex or TMT11plex Mass Tag Labelling Kit (ThermoFisher, Catalog Number A34808). TMT labelling efficiency was measured by LC-MS3 analysis after combining equal volumes (equivalent to ~ 1 μg each) from each sample. At this stage a ratio check was performed in which the total peptide intensities from each sample were compared for equivalence. Equal amounts of labelled peptide from each sample (as judged from ratio check data) were then combined for subsequent fractionation in a single HPLC run; each run involved a total amount of approximately 600 μg protein.

Quenching of TMT labelling reactions was performed by adding hydroxylamine to a final concentration of 0.5% (v/v) and incubating samples for 15 minutes at room temperature. Formic acid (FA) was added to a final volume of 2% (v/v) to lower the pH below 3.0 and samples were combined and de-salted using a SepPak tC18 Vac RC Cartridge (50 mg, Waters, Catalog Number WAT054960). HPLC fractionation was performed over a period of 75 minutes using an Agilent 1200 Series instrument with a flow rate of 600 μl/minute. Peptides were collected in a 96-well plate over a 65 min-gradient of 13-44%B with Buffer A comprising 5% acetonitrile, 10 mM ammonium bicarbonate, pH 8 and Buffer B comprising 90% acetonitrile, 10 mM ammonium bicarbonate, pH 8. Fractions were pooled to generate a total of 12-24 aliquots, followed by sample clean-up using the Stage Tip protocol with C18 Empore™ Extraction Disks (Fisher Scientific, Catalog Number 14-386-2). The matrix was primed with methanol and equilibrated with 70% acetonitrile, 1% FA followed by washing twice with 1% FA. Samples were loaded in 1% FA, followed again by two 1% FA washes, and finally peptides were eluted using 70% acetonitrile, 1% FA. Samples were dried before resuspension in MS Loading Buffer (3% acetonitrile, 5% FA).

#### LC-MS

The first half of the dataset (runs 1-4) was recorded after peptide separation on 100 μm columns packed with 1.8 μm C18 beads with a pore size of 12 nm (Sepax Technologies Inc.). The second half of the data (runs 5-8) was obtained after peptide separation on 75 μm columns packed with 2.6 μm Accucore beads (Thermo Fisher Scientific). Peptides were injected onto 30-40 cm, 100 and 75 μm (internal diameter) columns, respectively, and separated using an EASY-nLC 1200 HPLC (ThermoFisher Scientific). The flow rate was 450 nl/min for the 100 μm columns and 300 nl/min for the 75 μm columns with a gradient of 6-28%B over 170 minutes with Buffer A comprising 3% acetonitrile, 0.4% FA and Buffer B comprising 100% acetonitrile, 0.4% FA for the 100 μm columns and 5-35%B over 240 minutes with Buffer A comprising 0.125% FA and Buffer B comprising 95% acetonitrile, 0.125% FA for the 75 μm columns. The columns were heated to 60°C using a column heater (constructed in-house). Samples from the HPLC were injected into an Orbitrap Fusion Lumos Tribrid MS (ThermoFisher, Catalog Number FSN02-10000) using a multi-notch MS3 method^15,16^. MS scans were performed in the Orbitrap over a scan range of 400-1400 m/z with dynamic exclusion. Rapid rate (runs 1-4) and Turbo rate (runs 5-8) scans were performed in the Ion Trap with a collision energy of 35% and maximum injection times of 120 ms and 200 ms, respectively. TMT quantification was performed using SPS-MS3 in the Orbitrap with a scan range of 100-1000 m/z and an HCD collision energy of 55%. Orbitrap resolution was 50,000 (dimensionless units) with maximum injection times of 120 ms and 450 ms, respectively. MS isolation windows were varied depending on the charge state.

### Data Analysis

Mass spectrometric data (Thermo “.RAW” files) were converted to mzXML format, to correct monoisotopic m/z measurements, and to perform a post-search calibration. Peptide spectrum matches were assigned with a SEQUEST (v.28 (rev. 12), (c) 1998-2007 Molecular Biotechnology, Univ. of Washington, J.Eng/S.Morgan/J.Yates licensed to Thermo Fisher Scientific Inc.) based software against a size-sorted forward and reverse database of the human proteome (Uniprot 02/2014) with added common contaminant proteins. The database search included reversed protein sequences and known contaminants such as human keratins, which were excluded from subsequent analyses. Linear discriminant analysis was used to adjust PSM false discovery rate to (FDR) < 1% by applying the target-decoy database search strategy. Filtering was performed as described previously^16^. During peptide assignment for all data, oxidized methionine (+15.9949 Da) was searched dynamically. All peptide searches considered TMT modification (+229.1629 Da) on N-termini and lysine residues as static modifications. For protein identification and quantification, shared peptides were collapsed into the minimally sufficient number of proteins using rules of parsimony. Peptides with a total TMT value of >200 and an isolation specificity of > 0.7 were included in the final dataset.

## Data Records

MS proteomics Level 1 Data on peptides have been deposited to the ProteomeXchange Consortium via the PRIDE^19^ partner repository with the dataset identifiers PXD015542 and PXD017494.

Proteome datasets are also available on Synapse (ID syn7802481): https://www.synapse.org/#!Synapse:syn7802481; these data include Level 2 data on protein intensities (Synapse ID: syn20632472) and Level 3 data in which protein levels are normalized within and across runs. The datasets are also available for download via the HMS LINCS database (at https://lincs.hms.harvard.edu/db/datasets/20352/ and http://lincs.hms.harvard.edu/db/datasets/20370/)

## Technical Validation

### Mass Spectrometry instrument Quality Control

Quality control checks for mass spectrometry were incorporated at multiple points in the workflow. To test for efficient digestion of samples, defined as <15% of potential proteolysis sites uncleaved, a “digest check” was performed using LC-MS/MS as described in the methods section. TMT labelling efficiency aims for modification of >95% of available sites and was determined by LC-MS analysis via dynamic searches for N-terminal peptide modification by TMT. A “ratio check” was also performed using LC-MS3 to determine relative amounts of labelled peptides in each of the multiplexed samples, as described in the methods section. The purpose of the ratio check is to ensure equal amounts of peptide are pooled in the sample run through the mass spectrometer.

### Reproducibility of results

73 samples (60 unique breast cancer cell lines and 13 technical or biological replicates) distributed across three breast cancer clinical subtypes (**Figure 2A**) were randomly divided into 8 runs. Each run had one or more bridge samples that comprised a mixed sample derived from six cell lines (HCC1806, Hs578T, MCF7, MCF10A, MDAMB231, SKBR3). By analysing the same bridge sample in each MS run it was possible to compare runs to each other (see below). A total of five biological replicates of the MCF 10A cell line were also present in the eight batches as a further measure of data reproducibility.

**Figure 2.**
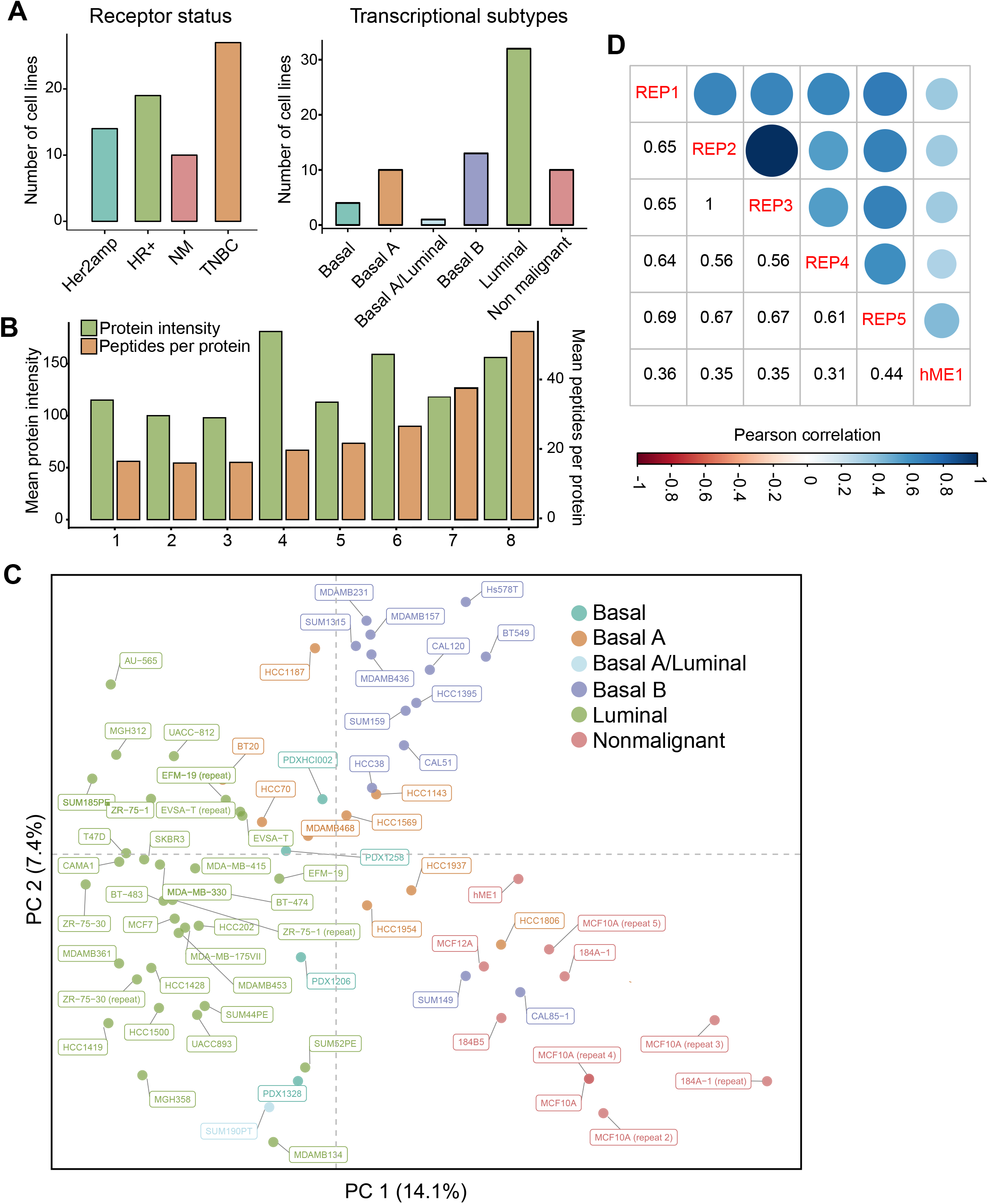
Distribution of samples for TMT LC/MS across cell lines and breast cancer subtypes. A total of 73 total samples were analyzed, representing 60 different cell lines and 13 technical and biological replicates for a subset of these lines. (A) Classification of samples based on molecular subtype (left panel) or receptor status (right panel). (B) Across-run-variability in mean protein intensities (green) and mean number of peptides reported per protein (orange) prior to normalization. (C) Principal Component Analysis (PCA) of the dataset following normalization to correct for run-specific differences in intensities and number of peptides quantified per protein. Clustering was based on known transcriptional subtypes. (D) Correlation between technical replicates of a single cell line (the non-malignant MCF 10A cell line) across five batches after normalization. HME1 is included to show contrast with another non-malignant cell line.

Principal component analysis (PCA) was performed prior to data normalization and revealed a significant degree of clustering by batch (runs 1-4 versus runs 5-8). This was true despite the high overlap in actual proteins detected. We could identify two reasons for this batch effect: advances in instrumentation and analytical methods meant that later batches exhibited better signal to noise (**Figure 2B,** upper panel), as assessed by an increase in mean intensity per protein. Samples in the second batch, had a greater number of quantifiable peptides per protein (**Figure 2B,** lower panel). A second difference between the two batches was in the number of aliquots generated by pooling HPLC fractions prior to MS. More aliquots were used in later MS runs (see the PRIDE^19^ datasets PXD015542 and PXD017494 for the number of runs per batch). To correct for these differences, the measured intensities for each protein in a run were normalized by the number of peptides detected for that protein. The samples in each run were then normalized to the bridge sample for that run so that the summed intensity scores across all samples were equivalent (within run normalization of samples). Finally, all samples were normalized to the data from the bridge sample of a reference run (run 4 in this study) to allow for comparisons across runs and batches.

### Biological validation

Subsequent to normalization, PCA based clustering of the 74 samples that were analysed by MS showed that data for each cell line clustered by transcriptional subtype (**Figure 2C**), suggesting that normalization was effective in removing batch effects. Further, the MCF10A replicates included in five runs comprising both batches clustered together, another indication that any remaining batch effects were small (**Figure 2D**). Out of 19,000 - 22,000 known human proteins^20^, we measured a total of ~13,000 unique proteins in our dataset. 7197 proteins were detected in all runs while the remaining proteins were observed to varying degrees in different runs. (**Figure 3A**). In shotgun proteomics, there is variation in the number of proteins detected in each MS run due to under-sampling and differences from one sample to the next can therefore reflect both real biological variation and under-sampling. Nonetheless, our normalized protein abundance data clustered the cell lines studied according to known subtype, *i.e* luminal, basal A, basal B and non-malignant, suggesting that protein features in the data capture the intrinsic heterogeneity of the cell line panel as previously established by transcript profiling (**Figure 2C**).

**Figure 3.**
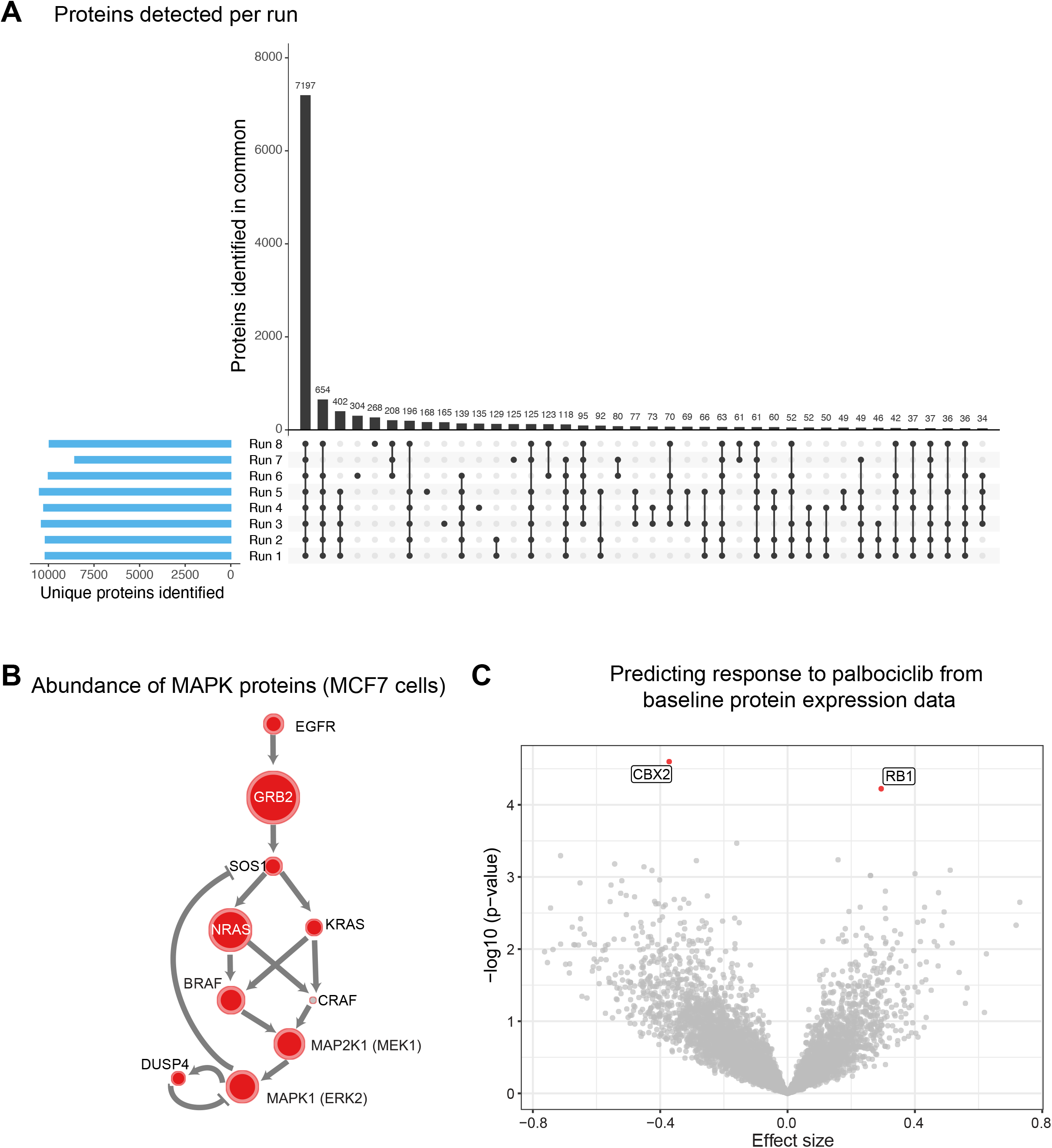
(A) UpSet plot to illustrate the depth in protein coverage for each of the eight batches and the extent of overlap in measured proteins across them. (B) Network representation of the MAPK signaling pathway with the size of the nodes (colored in red) scaled based on their measured abundance (log10 iBAQ) in MCF7 cells. The CRAF protein (colored grey) is included but was not detected in MCF7 cells, but was present in other lines. (C) Volcano plot (−log10 (*p*-value) against Effect size) shows the relative strength of each of the 7197 proteins in predicting breast cancer cell line response (GR AOC) to palbociclib. Negative effect size is associated with drug resistance and positive effect size is associated with sensitivity.

## Usage Notes

The NIH LINCS program has defined different data levels for all data, including proteomics, that comprise: Level 1 (primary or raw; .raw files and mzXML files in the case of MS data), Level 2 (relative peptide intensities reported for each run), Level 3 (sub-threshold peptides and contaminant proteins removed, and batch normalized) and Level 4 (signatures and markers of response)^21^ (these data levels are described in detail in the accompanying overview manuscript). Level 1 data for the current study are available for download from the Pride repository (PXD015542 and PXD017494). Level 2 data were generated using software provided by the Gygi Laboratory at HMS and comprise summed protein intensity estimates (available on Synapse: syn20632472). The level 2 data can also be generated from raw files using MaxQuant^22^. The protein intensity estimates in Level 2 data were then normalized using bridge samples to make cross-run and cross batch comparisons possible (available on Synapse: syn21585559). Protein expression in TMT MS data are relative estimates and can only be compared across samples on a protein-by-protein basis. Comparison of the absolute levels of different proteins is not possible because the TMT signal-to-noise ratio (s/n), which is used for relative quantification, depends on the injection time for each MS3 scan (variability in injection time means that relative, not absolute, s/n height is used to determine peptide quantity). Moreover, differences in peptide ionization, length and molecular weight as well as sampling biases that affect the spectral intensities of individual peptides complicate comparisons between proteins. However, these limitations can be, at least, partly overcome by using the intensity based absolute quantification (iBAQ) method that normalizes the intensities quantified for any give protein by the number of its theoretically observable peptides.^23^ This provides an estimate of absolute protein levels so that proteins in a sample can be compared with each other, but the units of the quantitation are still arbitrary (iBAQ normalized data comprise a set of LINCS Level 3 data, available on Synapse: syn21585566).

The intensity of the MS1 peak is proportional to the number of ionized peptide molecules at the measured mass to charge ratio (m/z) of a given analyte. Thus, absolute quantification of analytes can be derived from MS1 spectra by spiking-in references of the same peptides in known amounts. To estimate the true concentrations of a subset of proteins in our samples, the bridge sample was spiked with 48 analytes in the UPS2 standard (Sigma Aldrich) selected to span the dynamic range of known protein concentrations. The iBAQ-calibrated intensities of the bridge sample were then calibrated against the iBAQ values of the 48 known analytes to derive estimates of absolute protein abundance in molecules/pg of cell. Finally, we calculated the pg weight for a given number of cells per cell line model to derive the number of molecules for each protein per cell. An advantage of performing this absolute qualification step is that it makes it possible for independent datasets collected at different times and by different laboratories be integrated and compared. As an example, when we looked at the levels of proteins in the MAPK signalling pathway in MCF7 cells we found that SOS1 is an order of magnitude less abundant than other proteins in the pathway including GRB2, NRAS, and MAP2K1 (**Figure 3B**). This is consistent with previous data showing that SOS1 levels may be a limiting factor in receptor tyrosine kinase-mediated responses to growth factors^24^

Genomic and transcriptomic data have frequently been used to predict drug response and identify potential predictors or determinants of drug sensitivity^25–27^. As a first step in determining the utility of protein expression data in predicting drug response, we measured the responses of 56 breast cancer cell lines to the CDK4/6 inhibitor palbociclib (available on the LINCS database https://lincs.hms.harvard.edu/db/datasets/20343). Using iBAQ abundance for each of the 7197 proteins measured in all cell lines, we built univariate linear models to predict response (area over the GR curve) to palbociclib. The model included receptor status as a covariate to account for subtype specific differences in protein expression (The ‘lm’ package in R was used to encode the linear models using the formula “palbociclib GR AOC ~ protein expression + receptor status”) (**Figure 3C**). As expected, the abundance of RB1, a key substrate of CDK4/6 and mediator of cell cycle arrest^28^, was among the strongest predictors of response to palbociclib (*p*-value = 5.9e-06) and was positively correlated with increased sensitivity. In contrast, expression of CBX2 was correlated with resistance to palbociclib (*p*-value = 2.5e-06). Overexpression of CBX2, a protein involved in DNA damage repair and chromatin homeostasis^29,30^, has been associated with upregulation of genes involved in cell cycle progression and worse 5-year survival in breast cancer patients^31^. The association of CBX2 expression with resistance to palbociclib in cell lines provides a rationale for considering it as a potential biomarker in humans and a possible therapeutic target to overcome resistance to CDK4/6 inhibitors. This preliminary analysis suggests that baseline protein expression in untreated cell line models can be used to generate testable hypotheses about factors that influence drug sensitivity and resistance.

## Code Availability

Computational tools to process data and plot figures shown in the paper are available on https://github.com/labsyspharm/lincs_proteomics_data_descriptor and https://github.com/datarail/msda.

## Acknowledgements

The datasets featured in this paper were collected and analyzed as part of the “The Library of Integrated Network-Based Cellular Signatures” (LINCS) program and funded by the NIH Common Fund program (U54 grant HL127365). The datasets are currently available under a Creative Commons License CC BY 4.0. We thank S. Gygi and members of his laboratory for valuable experimental and technical MS advice and R. Eisert for help with sample preparation.

## Author contributions

CEM, KS, MH, and PKS designed and conceived the study. CEM designed the experiments and CEM, MC, BG, CV, RAE, MJB, GAB and MK performed the experiments. KS designed the computational analysis and tools for this study and KS, MKN performed the analysis. PKS oversaw the experimental and computational research. MK, MJB, MKN, PKS, CEM, and KS wrote the manuscript.

## Declaration of interests

KS is currently an employee of Bristol-Myers Squibb Inc., MH is an employee of Genentech, and RAE is an employee of Pfizer Inc. They declare no conflicts of interest.

## Supplementary Table

| ***Cell line*** | ***Growth media*** | ***Growth***<br>***conditions*** |
| --- | --- | --- |
| **184A1** | MEBM (CC-3150) + 1% FBS + 1% P/S | 37°C, 5% CO<sub>2</sub> |
| **184B5** | MEBM (CC-3150) + 1 ng/ml CT + 1% P/S | 37°C, 5% CO<sub>2</sub> |
| **AU-565** | RPMI-1640 + 10% FBS + 1% P/S | 37°C, 5% CO<sub>2</sub> |
| **BT-20** | EMEM + 10% FBS + 1% P/S | 37°C, 5% CO<sub>2</sub> |
| **BT-474** | RPMI-1640 + 10% FBS + 1% P/S | 37°C, 5% CO<sub>2</sub> |
| **BT-483** | RPMI-1640 + 20% FBS + 1% P/S, 0.01 mg/ml BI | 37°C, 5% CO<sub>2</sub> |
| **BT-549** | RPMI-1640 + 10% FBS + 1% P/S, 1 ug/ml IN | 37°C, 5% CO<sub>2</sub> |
| **CAL-120** | DMEM + 10% FBS + 1% P/S | 37°C, 5% CO<sub>2</sub> |
| **CAL-51** | DMEM + 20% FBS + 1% P/S | 37°C, 5% CO<sub>2</sub> |
| **CAL-85-1** | DMEM + 10% FBS + 1% P/S | 37°C, 5% CO<sub>2</sub> |
| **CAMA-1** | EMEM + 10% FBS + 1% P/S | 37°C, 5% CO<sub>2</sub> |
| **EFM-19** | RPMI-1640 + 10% FBS + 1% P/S | 37°C, 5% CO<sub>2</sub> |
| **EVSA-T** | EMEM + 10% FBS + 2 mM L-glutamine, 1% P/S | 37°C, 5% CO<sub>2</sub> |
| **HCC1143** | RPMI-1640 + 10% FBS + 1% P/S | 37°C, 5% CO<sub>2</sub> |
| **HCC1187** | RPMI-1640 + 10% FBS + 1% P/S | 37°C, 5% CO<sub>2</sub> |
| **HCC1395** | RPMI-1640 + 10% FBS + 1% P/S | 37°C, 5% CO<sub>2</sub> |
| **HCC1419** | RPMI-1640 + 10% FBS + 1% P/S | 37°C, 5% CO<sub>2</sub> |
| **HCC1428** | RPMI-1640 + 10% FBS + 1% P/S | 37°C, 5% CO<sub>2</sub> |
| **HCC1500** | RPMI-1640 + 10% FBS + 1% P/S | 37°C, 5% CO<sub>2</sub> |
| **HCC1569** | RPMI-1640 + 10% FBS + 1% P/S | 37°C, 5% CO<sub>2</sub> |
| **HCC1806** | RPMI-1640 + 10% FBS + 1% P/S | 37°C, 5% CO<sub>2</sub> |
| **HCC1937** | RPMI-1640 + 10% FBS + 1% P/S | 37°C, 5% CO<sub>2</sub> |
| **HCC1954** | RPMI-1640 + 10% FBS + 1% P/S | 37°C, 5% CO<sub>2</sub> |
| **HCC202** | RPMI-1640 + 10% FBS + 1% P/S | 37°C, 5% CO<sub>2</sub> |
| **HCC38** | RPMI-1640 + 10% FBS + 1% P/S | 37°C, 5% CO<sub>2</sub> |
| **HCC70** | RPMI-1640 + 10% FBS + 1% P/S | 37°C, 5% CO<sub>2</sub> |
| **hTERT-HME1** | MEMB + Lonza CC-3150 kit | 37°C, 5% CO<sub>2</sub> |
| **Hs 578T** | DMEM + 10% FBS + 1% P/S | 37°C, 5% CO<sub>2</sub> |
| **MCF 10A** | DMEM/F12 (1:1) + 5% HS + 1% P/S, 20 ng/ml<br>EGF, 0.5 mg/ml HC, 10 µg/ml IN, 100 ng/ml CT | 37°C, 5% CO<sub>2</sub> |
| **MCF12A** | DMEM/F12 + 5% HS + 1% P/S, 20 ng/ml EGF, 0.5 mg/ml HC, 10 µg/ml IN, 100 ng/ml CT | 37°C, 5% CO<sub>2</sub> |
| **MCF7** | DMEM + 10% FBS + 1% P/S | 37°C, 5% CO<sub>2</sub> |
| **MDA-MB-134-VI** | L-15 + 20% FBS + 1% P/S | 37°C, 0% CO<sub>2</sub> |
| **MDA-MB-157** | L-15 + 10% FBS + 1% P/S | 37°C, 0% CO<sub>2</sub> |
| **MDA-MB-175-VII** | L-15 + 10% FBS + 2mM L-glutamine, 1% P/S | 37°C, 0% CO<sub>2</sub> |
| **MDA-MB-231** | DMEM + 10% FBS + 1% P/S | 37°C, 5% CO<sub>2</sub> |
| **MDA-MB-330** | L-15 + 20% FBS + 2mM L-glutamine, 1% P/S + 30 ng/ml EGF + 0.016 mg/ml IN + 2mM glutathione | 37°C, 0% CO<sub>2</sub> |
| **MDA-MB-361** | L-15 + 20% FBS + 1% P/S | 37°C, 0% CO<sub>2</sub> |
| **MDA-MB-415** | L-15 + 15% FBS + 2mM L-glutamine, 1% P/S, 10 µg/ml IN | 37°C, 0% CO<sub>2</sub> |
| **MDA-MB-436** | L-15 + 10% FBS + 1% P/S, 10 µg/ml IN | 37°C, 0% CO<sub>2</sub> |
| **MDA-MB-453** | L-15 + 10% FBS + 1% P/S | 37°C, 0% CO<sub>2</sub> |
| **MDA-MB-468** | L-15 + 10% FBS + 1% P/S | 37°C, 0% CO<sub>2</sub> |
| **MGH312** | RPMI + 10% FBS + 1% P/S | 37°C, 5% CO<sub>2</sub> |
| **MGH358** | RPMI + 10% FBS + 1% P/S | 37°C, 5% CO<sub>2</sub> |
| **PDX1206** | DMEM/F12 (3:1) + 7.5% FBS + 1% P/S, 0.125 ng/ml EGF, 25 ng/ml HC, 5 µg/ml IN, 8.6 ng/ml CT, 5 uM Y-27632 | 37°C, 5% CO<sub>2</sub> |
| **PDX1258** | DMEM/F12 (3:1) + 7.5% FBS + 1% P/S, 0.125 ng/ml EGF, 25 ng/ml HC, 5 µg/ml IN, 8.6 ng/ml CT, 5 uM Y-27632 | 37°C, 5% CO<sub>2</sub> |
| **PDX1328** | DMEM/F12 (3:1) + 7.5% FBS + 1% P/S, 0.125<br>ng/ml EGF, 25 ng/ml HC, 5 µg/ml IN, 8.6 ng/ml CT,<br>5 uM Y-27632 | 37°C, 5% CO<sub>2</sub> |
| **PDXHCI002** | DMEM/F12 (3:1) + 7.5% FBS + 1% P/S, 0.125<br>ng/ml EGF, 25 ng/ml HC, 5 µg/ml IN, 8.6 ng/ml CT,<br>5 uM Y-27632 | 37°C, 5% CO<sub>2</sub> |
| **SK-BR-3** | McCoy's + 10% FBS + 1% P/S | 37°C, 5% CO<sub>2</sub> |
| **SUM1315MO2** | F-12 + 5% FBS + 1% P/S, 10 ng/ml EGF, 5 µg/ml<br>IN, 10 mM HEPES | 37°C, 5% CO<sub>2</sub> |
| **SUM149PT** | F-12 + 5% FBS + 1% P/S, 1 µg/ml HC, 5 µg/ml IN,<br>10 mM HEPES | 37°C, 5% CO<sub>2</sub> |
| **SUM159PT** | F-12 + 5% FBS + 1% P/S, 1 µg/ml HC, 5 µg/ml IN,<br>10 mM HEPES | 37°C, 5% CO<sub>2</sub> |
| **SUM-185PE** | Ham's F12 + 5% FBS + 1% P/S + 5 µg/ml BI + 1<br>µg/ml HC + 10 mM HEPES | 37°C, 5% CO<sub>2</sub> |
| **SUM-190PT** | Ham's F12 + 1% FBS + 1% P/S, 5 µg/ml BI, 1<br>µg/ml HC, 10 mM Hepes, 5 mM ethanolamine, 5<br>µg/ml transferrin, 10 nM triiodo thyronine, 50 nM<br>sodium selenite, 1 g/l BSA | 37°C, 5% CO<sub>2</sub> |
| **SUM-44PE** | Ham's F12 + 1% FBS + 1% P/S, 5 µg/ml BI, 1<br>µg/ml HC, 10 mM HEPES, 5 mM ethanolamine, 5<br>µg/ml transferrin, 10 nM triiodo thyronine, 50 nM<br>sodium selenite, 1 g/l BSA | 37°C, 5% CO<sub>2</sub> |
| **SUM-52PE** | Ham's F12 + 5% FBS + 1% P/S + 5 µg/ml BI + 1 µg/ml HC + 10 mM HEPES | 37°C, 5% CO<sub>2</sub> |
| **T47D** | RPMI-1640 + 10% FBS + 1% P/S, 1 µg/ml IN | 37°C, 5% CO<sub>2</sub> |
| **UACC-812** | L-15 + 20% FBS + 2mM L-glutamine, 1% P/S, 20 ng/ml EGF | 37°C, 0% CO<sub>2</sub> |
| **UACC-893** | L-15 + 10% FBS + 1% P/S | 37°C, 0% CO<sub>2</sub> |
| **ZR-75-1** | RPMI-1640 + 10% FBS + 1% P/S | 37°C, 5% CO<sub>2</sub> |
| **ZR-75-30** | RPMI-1640 + 10% FBS + 1% P/S | 37°C, 5% CO<sub>2</sub> |

Abbreviations: fetal bovine serum (FBS), penicillin/streptomycin (P/S), insulin (IN), bovine insulin (BI), hydrocortisone (HC), epidermal growth factor (EGF), cholera toxin (CT).

## Notes

https://lincs.hms.harvard.edu/db/datasets/20352/

### Competing Interest Statement

The authors have declared no competing interest.

